# Chronic ethanol self-administration alters dopamine in the caudate nucleus and putamen of rhesus macaques in a sex-dependent manner

**DOI:** 10.64898/2026.02.12.705407

**Authors:** Charles C. Levy, Verginia C. Cuzon Carlson, Kathleen A. Grant, Armando G. Salinas

## Abstract

Alcohol use disorder (AUD) affects over 28 million people in the U.S and is associated with neurobiological alterations, including in the basal ganglia. Within the basal ganglia, the caudate nucleus (caudate) and putamen are implicated in AUD due to their roles in ethanol reinforcement, with the caudate receiving inputs from cortico-associative areas and the putamen receiving inputs from somatosensory areas, supporting goal-directed and habitual behaviors respectively. These distinct behavioral roles are supported by dopamine signaling, including phasic dopamine, involved in assessing action-outcome associations, and tonic dopamine, which reflects ongoing dopaminergic tone that biases action initiation. Intrastriatal dopamine release is modulated by cholinergic interneurons via nicotinic acetylcholine receptors. Dysregulation of these mechanisms can contribute to the transition from occasional to habitual ethanol drinking. Here, we used in-vitro fast-scan cyclic voltammetry to measure dopamine signaling in male (n=6) and female (n=6) rhesus macaques following six months of ethanol self-administration. In putamen, ethanol increased tonic dopamine in both sexes, with females exhibiting greater release and faster dopamine uptake rates than males. In the caudate, ethanol self-administration enhanced dopamine uptake rates only in males. Phasic dopamine release was enhanced in caudate of both sexes but only putamen in males. nAChR blockade revealed that phasic dopamine release in males, but not females, was dependent on cholinergic modulation. These results demonstrate basal and sex-specific dopamine release and uptake are uniquely altered in rhesus macaque caudate and putamen in conjunction with chronic ethanol drinking.

## Introduction

The caudate nucleus (caudate) and putamen are implicated in alcohol use disorder (AUD) due to their distinct but complementary roles in ethanol consumption^1-9^. The caudate supports associative processes, whereas the putamen is critical for the formation of habitual behaviors ^10-15^. Dysregulation of these brain regions can lead to dysfunction characterized by the transition from initial, hedonic ethanol-seeking behavior to habitual or compulsive ethanol seeking as observed in AUD^2,16-18^. Both regions receive nigrostriatal dopamine (DA) projections, which become dysregulated following chronic ethanol exposure^19,20^. Previous literature indicates sex differences in dopamine function following chronic ethanol^21,22^, including in the caudate and putamen^23^. However, many of these studies did not directly compare male and female subjects. Here, we address this by using rhesus macaques that self-administered ethanol for over six months to determine how chronic ethanol consumption affects dopamine release dynamics in the caudate and putamen, with sex explicitly incorporated as a factor.

Dopamine neurons exhibit patterns of activity that vary across contexts and behavioral demands, supporting multiple aspects of motivated behavior^24^. Berke (2018) argues that dopamine signals are not simply defined as phasic dopamine equaling learning and tonic dopamine equaling motivation. According to Berke, phasic signaling is characterized by brief, high-frequency bursts that convey reward prediction errors (RPEs), which reflect the discrepancy between expected and actual outcomes^24^. Positive RPEs occur when outcomes exceed expectations, whereas negative RPEs occur when outcomes fall short^24^. Through reward error signaling, phasic dopamine updates action-outcome associations and could modulate goal-directed control of ethanol seeking, strengthening actions that yield unexpectedly positive outcomes and weakening those which do not.

Tonic dopamine signaling reflects the ongoing availability of dopamine within striatal circuits, modulating neural excitability and shaping how dopamine signals influence behavior^24^. Rather than encoding reward value, tonic dopamine biases action initiation, persistence and engagement^24-26^. Chronic ethanol exposure can disrupt both phasic and tonic release, producing region-specific alterations in dopamine release which could contribute to a shift from flexible, outcome-sensitive control toward habitual, maladaptive ethanol seeking^27,28^.

Local caudate and putamen circuitry strongly influence dopamine release. Cholinergic interneurons (CINs) within these regions modulate dopamine release via nicotinic acetylcholine receptors (nAChRs) on dopamine axons, providing a mechanism by which local circuits can dynamically regulate both phasic and tonic signaling^29-32^. Chronic ethanol disrupts this signaling^15,23,33-36^, potentially contributing to the shift from occasional to habitual ethanol-seeking behavior. Understanding how these local circuits adapt to prolonged ethanol exposure is critical for elucidating the neurobiological mechanisms underlying AUD.

Sex differences in dopamine signaling have been reported in both human^37,38^ and preclinical models^9,39-41^, yet many studies investigating ethanol-induced dopaminergic adaptations do not explicitly include sex as a biological variable^42^. Evidence suggests that males and females may differ in dopamine release^41,43-47^, dopamine uptake rates^40,41,43,48^, and responses to chronic ethanol^23,34^, highlighting the importance of examining these factors in studies of AUD^9^. Despite this, the sex-dependent effects of chronic ethanol on dopamine signaling in the caudate and putamen remain poorly characterized.

Using a translationally relevant nonhuman primate model of voluntary self-administration, we examined how chronic ethanol consumption affects dopamine release dynamics in the caudate and putamen. This model allows for species translational insight into the neural adaptations that contribute to the transition from associative processes to sensory motor processes, addressing gaps in the literature that have largely focused on male rodents. By integrating phasic and tonic dopamine release dynamics and CIN-mediated local control, this study provides a mechanistic perspective on how chronic ethanol reshapes striatal dopamine signaling to excessive ethanol consumption.

## Methods

### Subjects

Ex vivo tissue was requested through the Monkey Alcohol Tissue Research Resource (MATRR) from three cohorts of male and female rhesus macaques (Macaca mulatta) from the breeding colony at Oregon National Primate Research Center (ONPRC) and used for fast-scan cyclic voltammetry experiments. The rhesus cohort 18 (n=12; 6 males, 6 females; ages 6.4-6.9 years old at necropsy) served as the ethanol self-administration group while rhesus cohort 19 (n=8; 4 males, 4 females; ages 6.3-7.4 years at necropsy) and rhesus cohort 22 (n=3; 3 females; ages 4.5-5.4 years at necropsy) served as controls with maltodextrin substituted for ethanol. Rhesus macaques were individually housed for 22 hours and paired with a social partner for 1-2 hours per day under controlled environmental conditions (11-hour light cycle from 06:00 to 17:00; temperature maintained at 20-22 °C; humidity at 65%), beginning six months prior to the ethanol induction phase. They were fed a nutritionally complete diet in the form of 1g banana flavored pellets (Test Diet, Inc.) in a daily quantity that allow for weight gain throughout the experiment. Their diet was supplements with fresh fruit and vegetables daily and body weights were taken weekly. All procedures were conducted in accordance with the Guide for the Care and Use of Laboratory Animals and approved by ONPRC IACUC. All timeline, drinking data, and cohort details can be found on the MATRR website (www.matrr.com).

### Ethanol Self-Administration

Rhesus macaques underwent an established training and drinking protocol as previously described^49-52^, and the experimental timeline is depicted in Figure 1A. Using positive reinforcement, rhesus macaques were trained to present their legs for awake blood draw and to pull a dowel and finger poke to operate a panel. Pulling the dowel allowed access to water from one of two drinking spouts and banana flavored food pellets were delivered under a 5-minute fixed-time schedule. This schedule of food delivery is documented to induce polydipsia in macaque monkeys^53^. All monkeys underwent four months of schedule-induced polydipsia (SIP), in which required volumes of either water or 4% ethanol (w/v) needed to be consumed to escape the fixed schedule of individual pellets that were delivered on a fixed time of 5-minutes. For the first 30 days of SIP, monkeys had to consume daily a volume of water that was equivalent to a 1.5 g/kg dose of ethanol. Then every 30 days the dose of ethanol increased such that monkeys had to consume volumes of 4% ethanol that would be equivalent to a dose of 0.5 g/kg, 1.0 g/kg, and 1.5 g/kg. Following SIP, rhesus macaques entered approximately six-months (162-177 days) of an “open-access” phase with ad libitum access to the 4% ethanol solution and water for 22 hours daily. Based on consumption patterns during this period, rhesus macaques could be categorized into four groups: low drinkers (ethanol intake below binge levels), binge drinkers (intake > 2 g/kg on > 55% of days with blood ethanol concentrations (BECs) > 80 mg/dL), heavy drinkers (intake > 3 g/kg on > 20% of days), and very high drinkers (>10% of open access days with an ethanol intake exceeding 4 g/kg)^51^. BECs were determined from blood collections at 7 h into the 22 h session every 5-7 days.

**Fig. 1.**
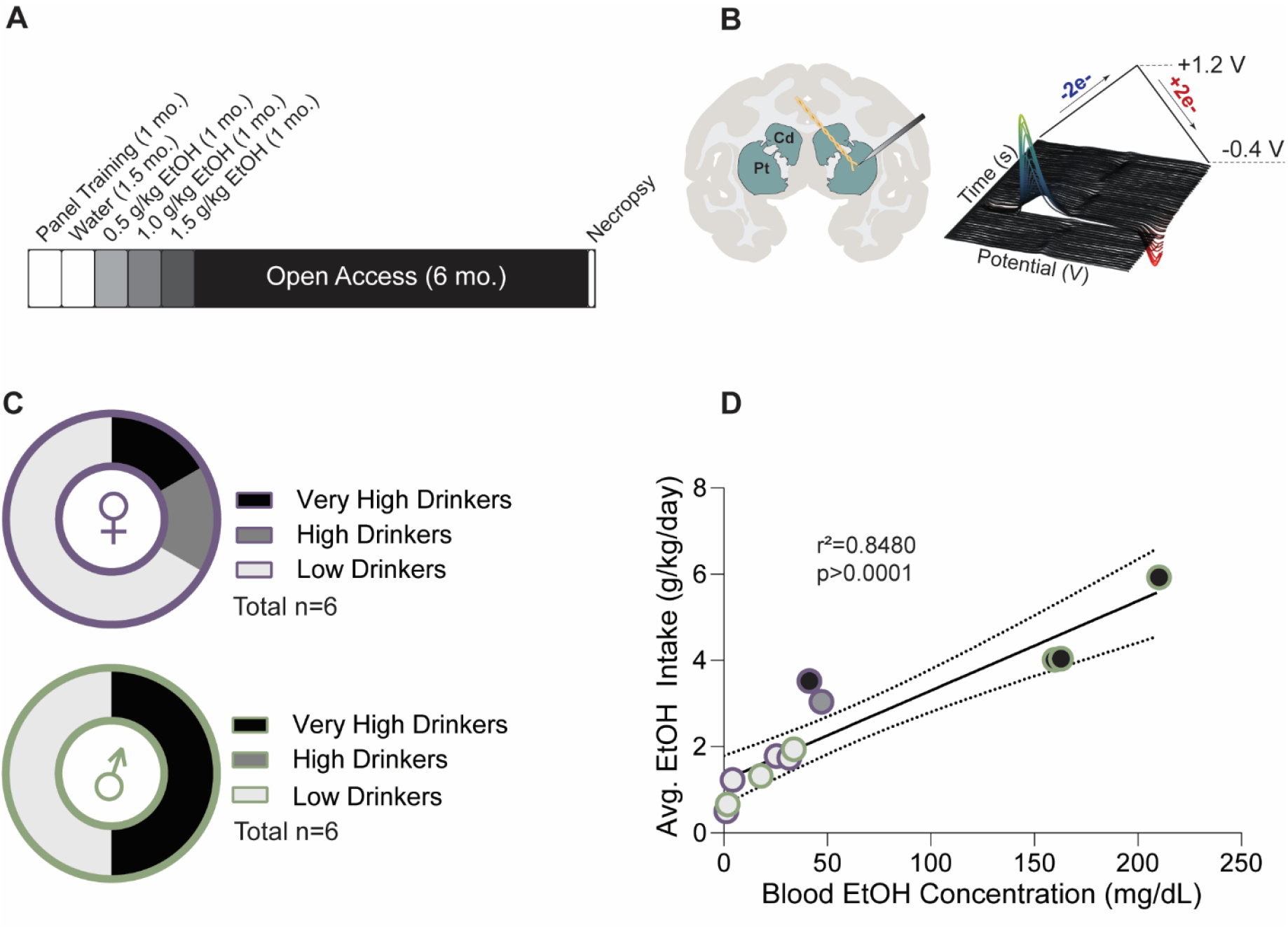
Experimental design and rhesus macaque ethanol consumption. **A** Rhesus macaque ethanol drinking paradigm. **B** Schematic of fast-scan cyclic voltammetry on rhesus macaque brain slice with caudate nucleus and putamen labeled. **C** Breakdown of ethanol drinking categories based on consumption patterns (low, high, very high dnnkers) and sex. **D** Average ethanol intake positively correlated with average blood ethanol concentration (BECs) (simple linear regression, r2 = 0.8480. p <0 0001) BECs were collected seven hours after onset of the session.

### Ex vivo brain slice preparation

Procedures for tissue collection and ex vivo brain slice preparation have been described in detail^23,54,55^. Briefly, rhesus macaques were anesthetized with sodium pentobarbital (30-50 mg/kg, i.v.) and transcardially perfused with a cold oxygenated perfusion solution (in mM: 124 NaCl, 23 NaHCO_3_, 3 NaH_2_PO_4_, 5 KCl, 2 MgSO_4_, 10 d-glucose, 2 CaCl_2_). Following perfusion, a craniotomy was performed, and the brain was extracted and blocked using a macaque coronal plane brain matrix (Electron Microscopy Sciences, Hatfield, PA). A tissue block containing the caudate and putamen from the left hemisphere was dissected and sectioned using a VT1200S vibratome (Leica, Buffalo Grove, IL) in cold, sucrose-based cutting solution (in mM: 194 sucrose, 30 NaCl, 4.5 KCl, 1 MgCl_2_, 26 NaHCO_3_, 1.2 NaH_2_PO_4_, 10 Glucose, pH 7.4), continuously bubbled with 95% O_2_/5% CO_2_. Coronal slices (250 µm thick) were cut using a ceramic blade (Camden Instruments Limited, Lafayette, IN). Slices were then transferred to voltammetry artificial cerebrospinal fluid (vACSF) (in mM: 126 NaCl, 2.5 KCl, 1.2 NaH_2_PO_4_, 2.4 CaCl_2_, 1.2 MgCl_2_, 25 NaHCO_3_, 11 glucose, 20 HEPES, 0.4L-ascorbate, pH 7.4) and incubated at 33 °C for 1 hour, followed by an additional hour at room temperature prior to the start of experiments.

### Fast Scan Cyclic Voltammetry

Carbon fiber electrodes (CFEs) were fabricated by inserting a single 7 µm diameter carbon fiber (T650 fiber; Goodfellow, Coraopolis, PA) into a 1.2 mm OD/0.68mm ID borosilicate glass capillary tube (A-M Systems, Carlsborg, WA). A tight seal around the fiber was achieved using a micropipette puller, and the exposed fiber was trimmed under high magnification to a final length of 150-250 µm beyond the glass seal.

Dopamine was measured by applying a triangular wave form (−1.2 V to +1.2 V to −1.2V; 400V/s; 10Hz) to the CFE using a Chem-Clamp potentiostat (Dagan Corporation, Minneapolis, MN) and DEMON Voltammetry and Analysis software as previously described^23,56^ (Figure 1B, right). To evoke dopamine release, a twisted stainless steel bipolar stimulating electrode was placed onto the tissue followed by placement of the CFE ∼300µm from the stimulating electrode (Figure 1B, left). Input-output curves were collected with single electrical pulses of varying intensities (200 µA, 400 µA, 800 µA, or 1600 µA) with a DS3 Constant Current Stimulator (Digitimer, Fort Lauderdale, FL). Each stimulus consisted of a 2 ms monophasic pulse, delivered every four-five minutes. Carbon fiber electrodes were calibrated to 1µM DA after all experiments were completed.

To examine differences between tonic and phasic dopamine release, electrical stimuli were applied as a single pulse or a burst of six pulses at 50Hz before and after bath application of the nicotinic acetylcholine receptor (nAChR) antagonist, dihydro-β-erythroidine hydrobromide (DHβE; 1uM). This allowed us to pharmacologically assess the nAChR cholinergic contribution to tonic and phasic DA release. Following 15 minutes of DHβE incubation, train stimulation protocols were applied. No more than two slices per rhesus macaque and brain region were included in analyses.

### Drugs

Dopamine-HCL was obtained from Sigma-Aldrich (St. Louis, MO) and dihydro-β-erythroidine hydrobromide was obtained from Tocris Bioscience (Minneapolis, MN). All drugs were dissolved into vACSF. All chemicals were obtained from Sigma Aldrich (St. Louis, MO).

### Data analysis & Statistics

Data was entered into excel and imported into GraphPad Prism 10 for statistics. For input-output curve comparison, three-way ANOVA (treatment, sex, stimulation-intensity) was used. For tau analysis, two-way ANOVA (sex and treatment) was used. For cholinergic contribution to dopamine release comparison, two-way ANOVA (sex and treatment) was used. For phasic dopamine enhancement comparison, two-way ANOVA (sex and treatment) was used. To determine if dopamine release was significantly increased relative to single pulse, responses were normalized to single pulse and compared to this value (100%) using a two-tailed one-sample t-test. For phasic dopamine release in the presence of DHβE comparison, three-way ANOVA (treatment, sex, DHβE) was used. Figures were made using Prism and Adobe Illustrator.

## Results

Following six months of drinking, the cohort consisted of seven low drinkers (4 females, 3 males), 1 high drinker (female), and 4 very high drinkers (1 female, 3 males; Figure 1C). The average daily ethanol consumption over the 6 months of open access positively correlated with BECs (simple linear regression, F(1,10) = 55.81, p <0.0001) (Fig. 1D).

In the caudate and putamen (Figure 2A, 2E), input-output curves were obtained by varying the stimulation intensity (200-1600 μA) and measuring the concentration of dopamine released similar to the representative traves in Figures 2B and 2F. Stimulation input-output curves in the caudate revealed a significant main effect of stimulation intensity (mixed-effects three-way ANOVA, F(1.30, 32.02) = 46.82, p < 0.0001) (Fig. 2C). No main effects of treatment (F(1, 25) = 3.01, p = 0.095) or sex (F(1, 25) = 1.00, p = 0.33) were observed (Fig. 2C). To assess the effects of chronic ethanol self-administration on dopamine uptake rates, we determined the decay constant tau. Tau analysis revealed significant main effects of treatment (Two-way ANOVA F(1, 14) = 5.22, p = 0.039) and sex (F(1, 14) = 4.87, p = 0.045) as well as an interaction between treatment and sex (F(1, 14) = 4.87, p = 0.046) (Fig. 2D). Post hoc analysis revealed that ethanol consuming male rhesus macaques had significantly lower tau than male controls (Uncorrected Fishers LSD, t = 2.69, p = 0.018), with no treatment effects across the females (Fig. 2D). Post hoc analysis also revealed that male rhesus macaques had higher tau relative to female control (Uncorrected Fishers LSD, t =3.54, p = 0.0033) (Fig. 2D).

**Fig. 2.**
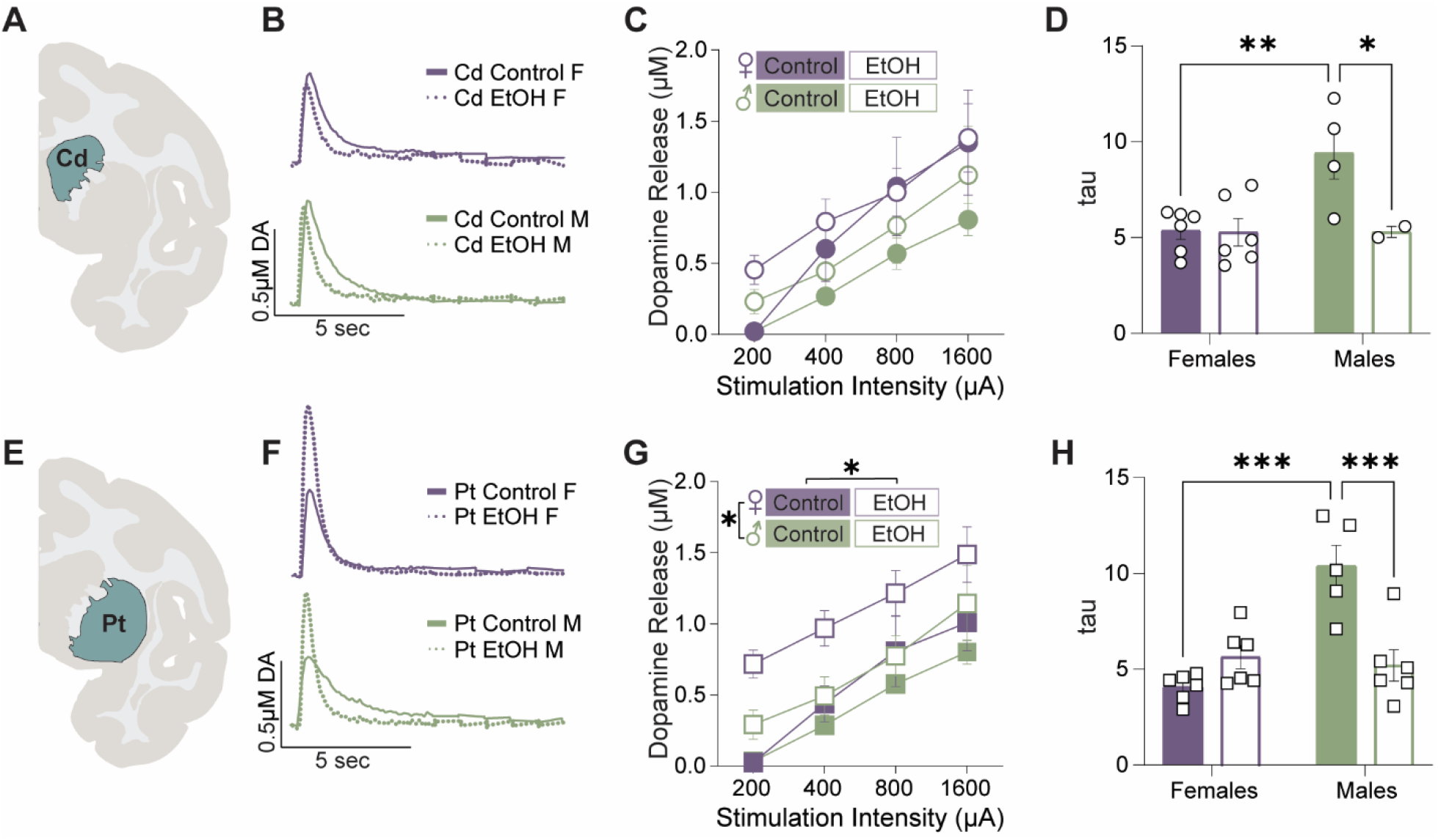
Chronic ethanol dysregulated dopam.ne release and uptake rates in a sex dependent manner across the caudate nucleus and putamen **A** Brain slice schematic with highlighted region of interest (caudate nucleus) for Fig. 2 B-D **B** Representative FSCV traces across treatment groups and sexes. **C** Chronic ethanol consumption did not alter evoked dopamine release in the caudate nucleus (n=5-8/group). **D** Chronic ethanol self-administration increased the uptake rate of dopamine in males, but not females. **E** Brain slice schematic with highlighted region of interest (putamen) for Fig 2 F-H. **F** Representative FSCV traces across treatment groups and sexes **G** In the putamen. chronic ethanol consumption increased evoked dopamine release in females, but not males (n=6-9/group) **H** Chronic ethanol self-administration increased the uptake rate of dopamine in males but not females * < 0.05, ^**^<001, ^***^<0 001

Stimulation input-output curves in the putamen revealed a significant main effect of stimulation intensity (mixed-effects three-way ANOVA, F(1.501, 38.03) = 98.73, p < 0.0001), treatment (mixed-effects Three-way ANOVA F(1, 26) = 6.42, p = 0.018), and sex (1, 26) = 10.42, p = 0.0034) (Fig. 2G). There were no significant interactions between sex, treatment, or stimulation intensity (all p > 0.2) (Fig. 2G). Similar to the caudate, tau analysis revealed significant main effects of treatment (Two-way ANOVA F(1, 19) = 6.14, p = 0.023), and sex (F(1, 19) = 16.28, p < 0.001) as well as an interaction between treatment and sex (F(1, 19) = 21.51, p < 0.001) (Fig. 2H). Post hoc analysis revealed that ethanol consuming male rhesus macaques had significantly lower tau than male controls (Uncorrected Fishers LSD, t = 4.915, p < 0.001), with no treatment effect among the females (Fig. 2H). Post hoc analysis also revealed that male rhesus macaques had higher tau relative to female control (Uncorrected Fishers LSD, t = 1.566, p < 0.001) (Fig. 2H).

To measure the cholinergic contribution to DA release, we measured DA release before and during bath application of 1μM DHβE (Figure 3A). Bath application of DHβE in the caudate (Figure 3B,C) revealed a significant main effect of sex where females had significantly less inhibition in the caudate compared to males (Two-way ANOVA F(1, 20) = 13.81, p = 0.0014), while treatment (F(1, 20) = 1.41, p = 0.25) and the interaction between sex and treatment (F(1, 19) = 0.04, p = 0.84) were not significant (Fig. 3D). Bath application of DHβE in the putamen (Figure 3E,F) revealed a main effect of sex where females had lower cholinergic contribution to dopamine release (Two-way ANOVA F(1, 20) = 13.81, p = 0.0014), while treatment (F(1, 20) = 1.41, p = 0.25) and the interaction between sex and treatment (F(1, 20) = 3.70, p = 0.070) was not significant (Fig. 3G).

**Fig. 3.**
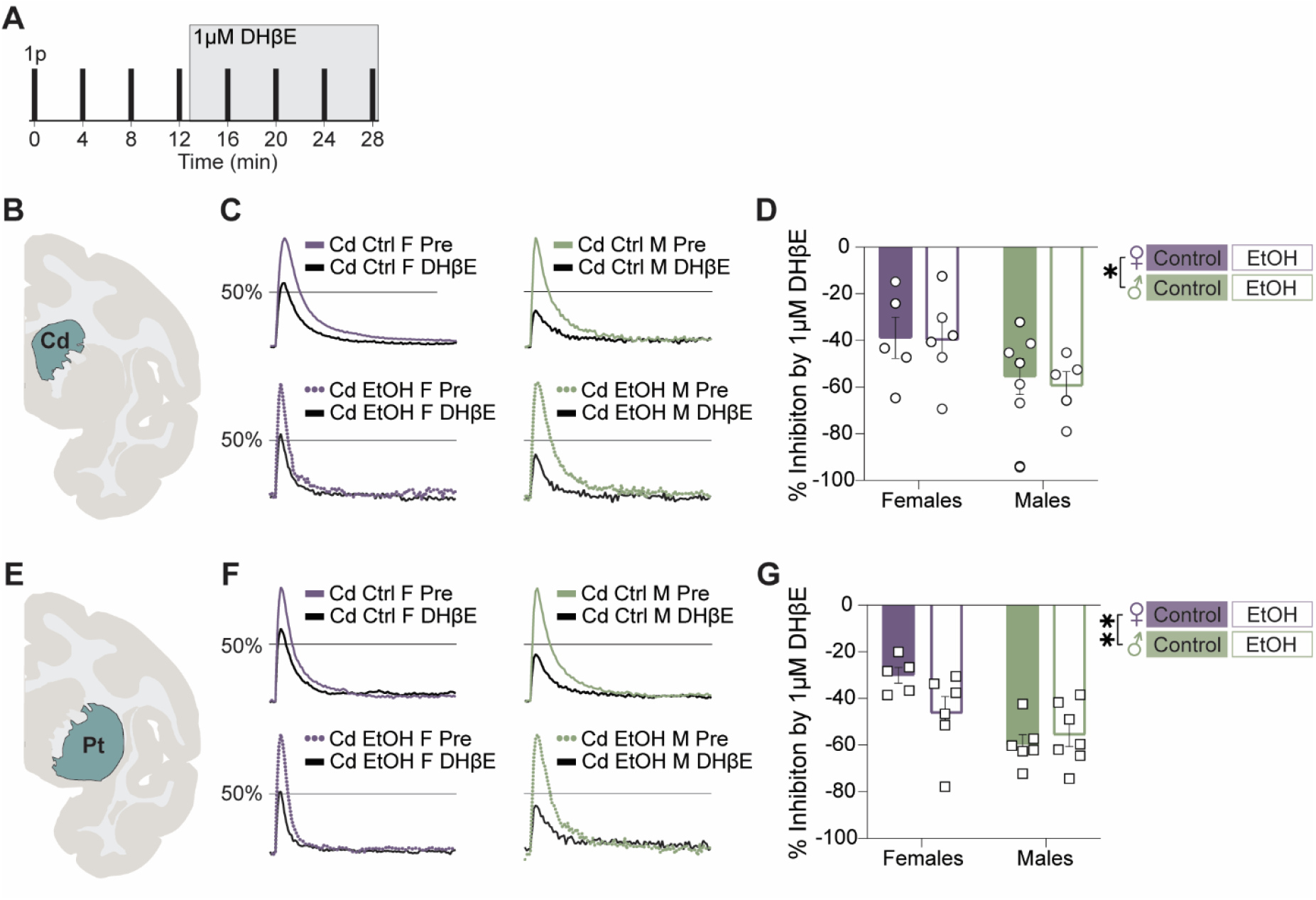
Cholinergic contribution to tonic dopamine release is lower in females than males in the caudate nucleus and putamen. **A** Schematic of experimental timeline of nicotinic acetylcholine receptor antagonist (DHβE) wash **B** Brain slice schematic with highlighted region of interest (caudate nucleus) for Fig. 3C-D. **C&D** Female rhesus macaques have less cholinergic contribution to tonic dopamine release compared to males in the caudate nucleus **E** Bram slice schematic with highlighted region of interest (putamen) for Fig. 3F-G **F&G** Female rhesus macaques have lower cholinergic contribution to tonic dopamine release compared to males in the putamen. ^*^ < 0.05, ^**^ < 0.01

To assess phasic DA release, we used a train consisting of 6 pulses applied at 50Hz (6p50Hz) and determined the percentage change from single-pulse-evoked responses (Figure 4A). In the caudate (Figure 4B), significant main effects of treatment (Two-way ANOVA F(1, 18) = 19.75, p = 0.0003) and sex (F(1, 18) = 6.96, p = 0.017) were observed (Fig. 4C). The interaction between sex and treatment (F(1, 18) = 0.058, p = 0.81) was not significant (Fig. 4C). Across all groups in the caudate, dopamine release was greater following the 6p50Hz train stimulation compared to the single pulse stimulation (One sample t-test Female control t(5) = 3.59, p = 0.016, Female ethanol t(4) = 8.12, p = 0.0013, Male control t(6) = 5.61, p = 0.0014, Male ethanol t(3) = 4.67, p = 0.019) (Fig. 4C). For the putamen (Figure 4E), a main effect of treatment (Two-way ANOVA F(1, 18) = 6.87, p = 0.017) was observed, while sex (F(1, 18) = 0.96, p = 0.34) and the interaction between sex and treatment (F(1, 18) = 1.02, p = 0.33) were not significant (Fig. 4F). Across all groups in the putamen, dopamine release was greater following the 6p50Hz train stimulation compared to the single pulse stimulation (One sample t-test Female control t(3) = 7.48, p = 0.049, Female ethanol t(4) = 3.74, p = 0.020, Male control t(5) = 4.31, p = 0.0076, Male ethanol t(6) = 3.77, p = 0.0093) (Fig. 4F).

**Fig. 4.**
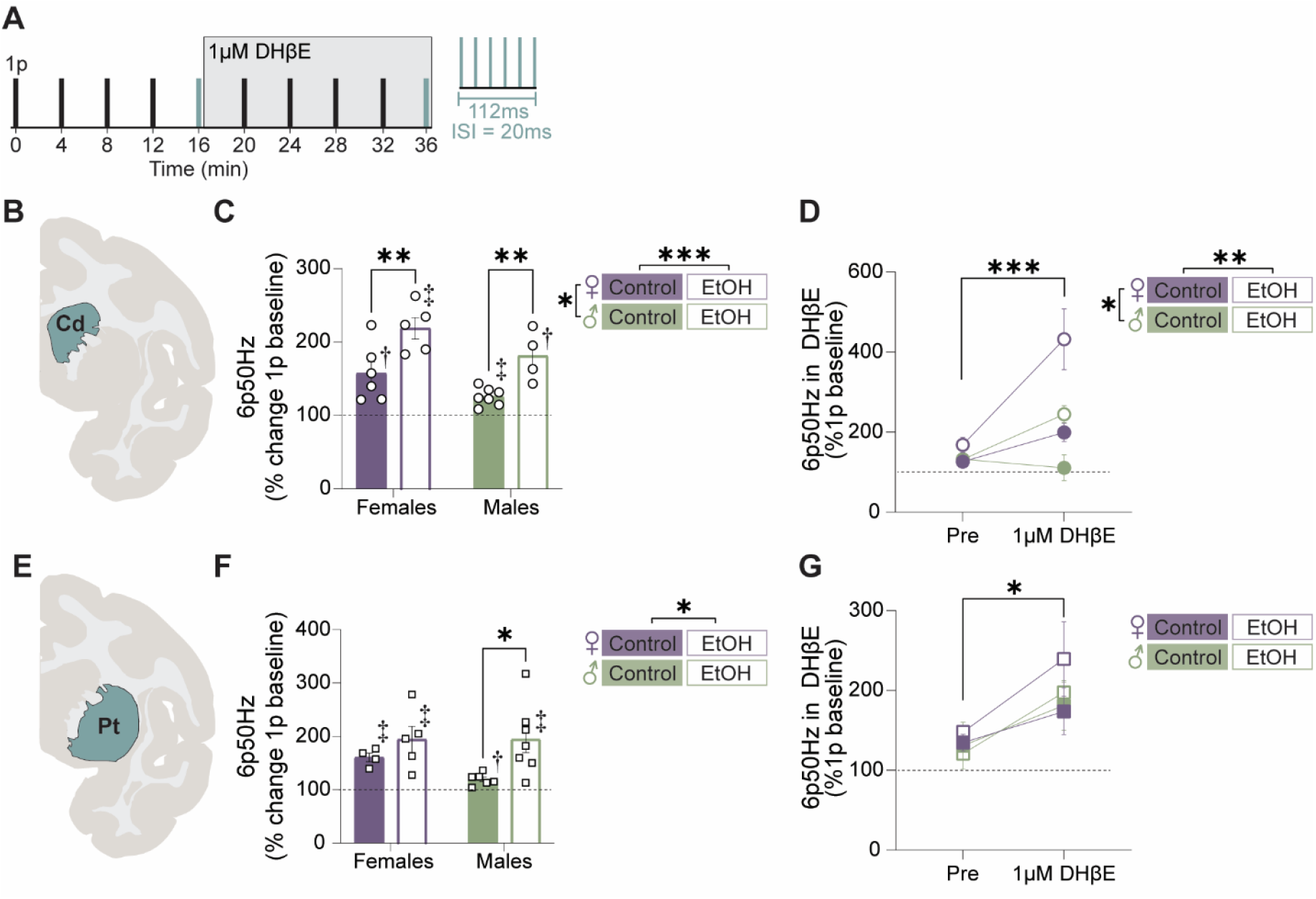
Chronic ethanol self-administration increases phasic dopamine release. A Schematic of experimental timeline of nicotinic acetylcholine receptor antagonist (DHl3E) application and 6p50Hz stimulations. **B** Brain slice schematic with highlighted region of interest (caudate nucleus) for Fig. 4 C-D. **C** Chronic ethanol self-administration increased phasic dopamine release in the caudate nucleus of both sexes. **D** Females have higher phasic dopamine release in the presence of DHl3E compared to males. **E** Brain slice schematic with highlighted region of interest (putamen) for Fig. 4 F-G. **F** Chronic ethanol self-administration increased phasic dopamine release in the putamen of males, but not females. **G** Phasic dopamine release in the presence of DHl3E compared to males. ^*^ < 0.05, ^**^ < 0.01, ^***^ < 0.001, < 0.05, < 0.01

We then assessed phasic dopamine release in the presence of DHβE to isolate dopamine terminals from the cholinergic contribution to DA release (Figure 4A, grey shaded box). In the caudate, significant main effects of DHβE, (Three-way ANOVA F(1, 12) = 26.93, p = 0.0002), treatment (F(1, 12) = 7.44, p = 0.018), and sex (F(1, 12) = 13.15, p = 0.0035) were observed (Fig. 4D). Whilst the interaction between sex and treatment (F(1, 12) = 1.58, p = 0.23) was not significant, a significant interaction between DHβE and treatment (F(1, 12) = 8.78, p = 0.012), as well as DHβE and sex (F(1, 12) = 15.49, p = 0.0020) was observed (Fig. 4D). There was no significant interaction between DHβE, sex, and treatment (F(1,12) = 0.47, p = 0.51 (Fig. 4D). In the putamen, a significant main effect of DHβE was observed (Three-way ANOVA F(1, 11) = 13.40, p = 0.0038) (Fig. 4G). There was no main effect of treatment (F(1, 11) = 0.72, p = 0.42) or sex (F(1, 11) = 1.25, p = 0.28) (Fig. 4G). There were no significant interactions observed (all p > 0.2) (Fig. 4G).

## Discussion

### Overview of Key Findings

Our results demonstrate that six months of voluntary ethanol self-administration produces sex- and region-specific alterations in dopamine signaling in the caudate and putamen of rhesus macaques. In the caudate, ethanol consumption altered dopamine uptake rates in a sex-dependent manner, with enhanced uptake rates selectively observed in ethanol-consuming males. In the putamen, ethanol increased tonic dopamine release in both sexes, alongside basal sex differences characterized by more dopamine release and faster uptake rates in females. Further, ethanol self-administration increased phasic dopamine release in the caudate of both sexes, and in the putamen in males. Lastly, we observed sex differences in the cholinergic modulation of dopamine release where females had less cholinergic contribution to dopamine release. Together, these findings indicate that chronic ethanol consumption results in sex-dependent adaptations in dopamine signaling which vary across the caudate and putamen.

### Basal Sex Differences in Dopamine Signaling

Previous rhesus macaque studies have not directly assessed sex effects, as cohorts consisted of single sexes and were often separated by a year. In contrast, male and female subjects within a cohort used in the present study were run in parallel. Notably, we employed two control cohorts’ separated by two years and found no differences in dopamine release and uptake, suggesting stable cohort responses. Females exhibited higher tonic dopamine release and faster uptake rates in the putamen, and faster uptake rates in the caudate compared to males. Sex differences in the putamen are consistent with prior reports^40,41,43,57^ and may reflect higher dopamine transporter expression and estradiol-mediated regulation^22,43,58^, both of which can increase extracellular dopamine availability^59^ and enhance responsiveness to rewarding or salient stimuli^9^.

In contrast, our evoked dopamine release measures in the caudate diverge from the rodent literature, which has generally reported higher extracellular striatal dopamine in females than males^40,41,43^. However, findings in humans have been mixed. For example, Munro et al., 2006 found that men had greater DA release than women. Yet, Manza et al., 2022 found no difference^60^. Additionally, in the present study females also exhibited less cholinergic modulation of dopamine release, suggesting that local circuit regulation differs by sex. Together, these intrinsic sex differences provide a framework for understanding region- and sex-specific effects of chronic ethanol exposure.

### Caudate Nucleus: Tonic Dopamine and Ethanol effects

Dopamine uptake rates in the caudate revealed sex-specific effects of ethanol. Under control conditions, females exhibited faster uptake rates than males. Following six months of ethanol self-administration, uptake rates were selectively increased in males. In contrast, tonic dopamine release in the caudate remained unchanged in both sexes, indicating that six months of daily drinking did not substantially alter dopaminergic tone in the caudate.

These baseline sex differences in uptake provide important context for interpreting ethanol-induced adaptations. Prior work has demonstrated reduced tonic dopamine following intermittent abstinence in a longer ethanol exposure paradigm (over 18 months)^23^. Given the lack of effect with six months of drinking, these data suggest that tonic signaling in the caudate is relatively resistant to continuous, short term ethanol self-administration. Consistent with this idea, males with one year of continuous, ethanol self-administration showed no change in tonic dopamine release. Only inclusion of intermittent abstinence periods reduced tonic dopamine in males^23^. In contrast to the present data, Siciliano et al., 2015 reported decreased tonic dopamine in the dorsolateral caudate of cynomolgus macaques following six months of ethanol self-administration using the same protocol. Discrepancies across studies may reflect differences in drinking levels, species or recording locations, as our study sampled randomly across the caudate^61^. Together, these findings suggest that tonic dopamine release in the caudate is largely preserved following relatively short-term ethanol self-administration.

### Putamen: Tonic Dopamine and Ethanol Effects

Dopamine uptake rates in the putamen revealed sex-specific effects of ethanol. Under control conditions, females exhibited faster uptake rates than males. Following six months of ethanol self-administration, uptake rates increased selectively in males. In contrast, tonic dopamine release in the putamen was enhanced by ethanol in both sexes, indicating that short-term ethanol exposure can elevate dopaminergic tone in the putamen. These intrinsic differences likely reflect sex-dependent regulation of dopamine signaling, including basal release and dopamine transporter function, and provide an important context for interpreting ethanol-induced adaptations.

This contrasts with our previous rhesus macaque study, which found that twelve months of ethanol self-administration increased dopamine uptake rates in females but not males^23^. These discrepancies suggest that sex-dependent adaptations in dopamine clearance are sensitive to the duration and pattern (i.e. intermittent abstinence) of ethanol consumption. These results are consistent with rodent work^41^ and reinforce the putamen’s role in the transition from associative processes to sensory motor processes in maintaining ethanol consumption. Additionally, our findings align with reports in rodents that females have higher tonic dopamine release and faster uptake rates in the striatum, though human studies have yielded mixed results.

### Phasic Dopamine Adaptations

Phasic dopamine signaling encodes RPEs and supports associative learning^24,62^. In the caudate, phasic dopamine release increased in both sexes following ethanol exposure, despite unchanged tonic dopamine availability. This enhancement may increase the salience of ethanol-associated cues and strengthen action-outcome associations, biasing behavior toward ethanol reinforcement. Importantly, baseline sex differences in tonic dopamine uptake create distinct circuit contexts. The faster uptake rates in females likely stabilize tonic dopamine availability, maintaining consistent engagement and action initiation, whereas males’ slower baseline uptake and ethanol-induced increases provide a less stable tonic background in which phasic RPE bursts may have a greater impact on updating action-outcome associations.

In the putamen, phasic dopamine release increased only in males, highlighting a region- and sex-specific adaptation. Since the putamen is critical for habit formation^12^, male-specific enhancement may accelerate the shift from more associative to more sensory driven mechanisms in chronic ethanol consumption. In this context, females’ higher baseline tonic dopamine supports sustained engagement and flexible, outcome-sensitive ethanol-drinking patterns, whereas males’ lower tonic dopamine combined with phasic enhancement increases the relative impact of RPE signaling on stimulus-driven behavior, potentially facilitating habit formation. Females showed no increase, indicating preserved flexible action-outcome control under short-term exposure.

### Phasic Dopamine and Sex-Specific Cholinergic Modulation

When phasic dopamine was measured in the presence of DHβE, which blocks nicotinic cholinergic input, we observed clear region- and sex-specific effects. In the caudate, females showed enhanced phasic release following ethanol exposure, suggesting that chronic ethanol induces terminal-level adaptations that increase responsiveness independent of local cholinergic modulation. In contrast, phasic release in males was unchanged.

In the putamen, blocking cholinergic input increased phasic release in both sexes. This finding suggests that the male-specific ethanol enhancement of phasic release observed under normal conditions is dependent on intact cholinergic signaling (Fig. 4F). Functionally, these results could suggest that females rely more on intrinsic dopamine terminal adaptations to maintain robust RPE signaling, whereas in males, phasic dopamine responses in putamen are more circuit dependent, relying on local cholinergic modulation.

## Conclusion

Together these findings demonstrate that six months of ethanol self-administration is sufficient to induce sex- and region-specific alterations in dopamine release dynamics within the caudate and putamen of rhesus macaques. In females, ethanol promotes terminal-level phasic enhancements in the caudate that are independent of cholinergic input, supporting stable RPE encoding and sustained goal-directed behavior. In males, ethanol-induced phasic adaptations are more dependent on local cholinergic inputs, especially in the putamen, which may accelerate the transition from cortico-associative driven flexible behaviors to more sensorimotor habitual-ethanol consumption. Discrepancy with prior non-human primate and rodent studies underscores the sensitivity of dopamine adaptations to sex and the duration of ethanol exposure. By utilizing a translationally relevant non-human primate model with paralleled assessment of males and females, this work addresses a critical gap in the literature regarding sex-dependent effects of chronic ethanol on dopamine signaling. Future studies will be needed to determine the mechanisms underlying these functional changes, including sex-specific alterations in dopamine transporter & nicotinic modulation of DA release expression.

## Acknowledgements

We are grateful to the Cuzon Carlson and Grant laboratories for their technical assistance and for hosting us while completing these studies. We would also thank Logan Slade and Morgan Schichtel for reviewing the manuscript and Anthony Huberty for assistance with schematics. This work was supported by National Institutes of Health grants R00AA025991 and R03AA030400 (AGS), P51OD011092 (KAG, VCC), P60AA010760 (KAG, VCC), U01AA013510 (KAG), U24AA013641 (KAG), and R24AA016431 (KAG, VCC).

The authors declare no competing interests.

## Contributions

CCL: Collected data, data analysis, manuscript preparation, manuscript editing VCC: Designed research, manuscript editing

KAG: Designed research, manuscript editing

AGS: Designed research, collected data, data analysis, manuscript preparation, manuscript editing, acquired funding

## Notes

### Competing Interest Statement

The authors have declared no competing interest.

